# Neuropathic pain and distinct CASPR2 autoantibody IgG subclasses drive neuronal hyperexcitability

**DOI:** 10.1101/2024.09.04.611282

**Authors:** Margarita Habib, Anna-Lena Wiessler, Patrik Greguletz, Michéle Niesner, Mareike Selcho, Ligia Abrante, Christian Werner, Annemarie Sodmann, Maximilian Koch, Zare Abdelhossein, Harald Prüss, Justina Dargvainiene, Jan Lewerenz, Robert Handreka, Péter Körtvélyessy, Dirk Reinhold, Franziska Thaler, Kalliopi Pitarokoili, Robert J. Kittel, Michael Briese, Michael Sendtner, Heike Rittner, Frank Leypoldt, Claudia Sommer, Robert Blum, Kathrin Doppler, Carmen Villmann

**Affiliations:** Institute for Clinical Neurobiology, University of Wuerzburg, Germany; Department of Neurology, University Hospital Wuerzburg, Germany; Department of Animal Physiology, Institute of Biology, Leipzig University, Germany; Institute of Clinical Chemistry, University Hospital Schleswig-Holstein Kiel/Lübeck, Kiel, Germany; Department of Biotechnology and Biophysics, Biocenter, Julius-Maximilians-University Wuerzburg, Germany; Department of Neurology and Experimental Neurology, Charité Universitätsmedizin Berlin, Germany; German Center for Neurodegenerative Diseases (DZNE) Berlin, Germany; Department of Neurology, University Hospital Ulm, Germany; Department of Neurology, Medical University Lausitz Carl Thiem, Cottbus, Germany; Labor Berlin, Innovations, Berlin, Germany; Institute for Molecular and Clinical Immunology, University Magdeburg, Germany; Institute of Clinical Neuroimmunology, University Hospital and Biomedical Center Ludwig-Maximilians University Munich, Germany; Department of Neurology, St. Josef Hospital, Ruhr-University Bochum, Germany; Department of Anesthesiology, Intensive Care, Emergency and Pain Medicine, Centre for Interdisciplinary Pain Medicine, University Hospital Wuerzburg, Germany; Department of Neurology, University Hospital Schleswig-Holstein, Kiel, Germany

**Keywords:** CASPR2 autoantibodies, neuropathic pain, IgG subclass, Kv potassium channels, hyperexcitability

## Abstract

Patients with autoantibodies (aAbs) against the contactin-associated protein-like 2 (CASPR2) suffer from a variety of clinical syndromes including neuropathic pain, in some patients even as the only symptom. CASPR2 is an adhesion protein of the neurexin IV family and part of the voltage-gated potassium channel complex (VGKC) in neurons of dorsal root ganglia (DRG). The subsequent pathological mechanisms following the binding of CASPR2 aAbs and their association with pain are only partially understood.

CASPR2 aAbs are mainly of the IgG4 subclass. Previous studies have neglected subclass-dependent effects. Here we investigated 49 subclassified patient serum samples positive for CASPR2 aAbs. To unravel underlying molecular mechanisms, we used a combination of super-resolution lattice structural illumination microscopy (SIM^2^) and functional readouts by calcium imaging and electrophysiological recordings.

CASPR2-positive patient sera subclassified in IgG4 together with at least one other IgG subclass (IgGX) and patients with only IgG4 were further subdivided into the pain and no pain group. Patient subclassification shed further light on the pathological mechanisms of CASPR2 aAbs. A decrease of CASPR2 expression after long-term exposure to CASPR2 aAbs was only observed for the patient group without pain. Upon withdrawal of the CASPR2 aAbs, CASPR2 expression returned to normal level. Structural alterations were obtained by increased distances between CASPR2 and associated potassium channels along DRG axons using high-resolution lattice SIM^2^ microscopy but only following binding of CASPR2 aAbs from patients with pain.

Similarly, CASPR2 aAbs of patients with pain significantly increased overall neuronal excitability of cultured DRG neurons as measured by calcium imaging. Patch-clamp recordings revealed significantly decreased current amplitudes of voltage-gated potassium (Kv) channels after incubation with all four CASPR2 aAbs subclassifications with the most prominent effect of serum samples harboring IgG4 aAbs. Notably, a patient serum sample lacking IgG4 did not alter Kv channel function. Withdrawal of aAbs rescued Kv channel function to normal levels suggesting that the affected potassium channel function is rather due to a functional block of the VGKC rather than altered structural integrity of the VGKC.

Taken together, we found IgG4 aAbs to be a major modifier of potassium channel function. The increase in DRG excitability is primarily due to impaired Kv channel conductance as a consequence of CASPR2 aAbs binding but additional and so far unidentified signal pathways contribute to this process in patients with neuropathic pain.

## Introduction

Autoantibodies (aAbs) against contactin-associated protein-like 2 (CASPR2) are associated with various clinical syndromes including limbic encephalitis, Morvan’s syndrome, peripheral nerve hyperexcitability syndrome, ataxia, pain, and sleep disorders.^1–3^ CASPR2 aAb-mediated diseases often exhibit clinical relapses (25%) mostly observed if immunotherapy is discontinued.^4,5^

A significant number of patients positive for CASPR2 aAbs suffer from neuropathic pain, and in some patients, this is the only symptom present.^6,7^ In our patient cohort (total of 115 patients), pain was a frequent symptom in 36% of all patients, often severe (64% of the patients with pain) and/or even the major symptom (54% of the patients with pain). Besides pain severity, two major phenotypes were identified (i) primarily distal-symmetric burning pain and (ii) widespread pain with myalgia and cramps.^8^ However, the mechanism by which CASPR2 aAbs drive neuropathic pain is poorly understood.

CASPR2 is an adhesion protein of the neurexin IV family organizing the voltage-gated potassium channel complex (VGKC) in the CNS and at juxtaparanodal sites in the PNS. The VGKC is formed by an interaction of CASPR2 to the intracellular protein 4.1b which is linked to the postsynaptic density protein 95 (PSD-95) which forms protein-protein interactions with potassium channels of the Kv1.1/Kv1.2 subtype.^9–12^ The proper clustering of Kv channels is an essential factor to keep the neuronal electrical properties enabling signal transmission.^11,13^ A direct mechanism by which CASPR2 organizes Kv channel clusters is not known.

It has been suggested that upon binding of CASPR2 aAbs, the associated potassium channels of the Kv subtype are altered in their somatic membrane expression which causes hyperexcitability of neurons and thus mediates neuropathic pain.^13^ Conversely, a significant increase in Kv1.2 expression upon presence of CASPR2 aAbs has been observed in transfected HEK-293 cells and hippocampal neurons.^11^

The CASPR2 aAbs are mainly of the IgG4 subclass.^5,14,15^ IgG4 differs from IgG1-3 by its inability to crosslink proteins and to induce subsequent protein internalization. Moreover, IgG4 is incapable to activate the complement system. This aligns with unaltered surface expression of CASPR2 or minor trends to enhanced expression of CASPR2 *in vitro* in hippocampal cultures and transfected HEK-293 cells or *in vivo* in mice injected with patient anti-CASPR2 IgG.^15,16^

The architecture of CASPR2 consists of a large extracellular domain with eight structural subdomains including four laminin G (L1-L4) domains, two epidermal growth factor (EGF)-like domains, a discoidin domain, and a fibrinogen-like domain, one transmembrane domain and a short intracellular C-terminal domain with the binding motif for the 4.1b protein. Twelve potential glycosylation sites are distributed at the extracellular N-terminal domain.^17–20^ A crystal structure is also only known for the extracellular domain of CASPR2 confirming the domain organization.^21^ The discoidin domain has been determined to be unique compared to other neurexin family members and contains the primary epitope for CASPR2 aAbs.^18,22^ Dorsal root ganglia (DRG) contain sensory neurons and have been determined as active mediators in the development of neuropathic pain. Moreover, they present a robust target for neuromodulation^23,24^ and may also be a target for nanotechnological analgesic treatment.^25^ DRG neurons express the proteins of the VGKC.^26^ CASPR2 aAbs have been shown to bind to DRG neurons resulting in decreased expression and function of the associated potassium channels thereby generating hyperexcitable DRGs. The affected potassium channels have been suggested to harbor the Kv1.1 subunit determined by blocking with dendrotoxin (DTX).^13^ However, subsets of DRG neurons differ in their Kv channel (Kv1-12) expression.^27^ To discriminate Kv channel compositions, DTX and conotoxin κM-RIIIJ are potent *in vitro* blockers for Kv1 channels.^28,29^

In this study, we investigated the functional mechanisms of anti-CASPR2 aAbs from patients with and without pain on nociceptive neurons and the role of IgG subclasses. By assessing the effect of anti-CASPR2 IgG on protein surface expression, assembly of the VGKC, the function of Kv channels and neuronal excitability, we found clear differences between different IgG subclasses of aAbs and sera of patients with and without pain.

## Materials and methods

### Patients

Sera of 17 patients with anti-CASPR2 aAbs were included in the present molecular study. Those sera were selected according to clear IgG subclassification and corresponding pain phenotype and availability of sufficient patient serum for the analysis. Patients (total 115; 102 with clinical data; 49 sera with subclass analysis with 40 sera subclassified and published in a recent study^8^) were prospectively recruited at the University Hospital Würzburg, Department of Neurology, after having given written and oral informed consent or were recruited via the German Network for Research on Autoimmune Encephalitis (GENERATE). Clinical and demographic data and aAb titer from the 17 patient sera (four patient groups: with and without pain and further discrimination to IgG4 or IgG4 and at least one additional IgG subclass with 3-5 patient samples each) investigated in this study were taken from the patients’ records or the GENERATE registry and are summarized in **Table 1**. Sera of four controls without any neurological symptoms or pain conditions were also included.

**Table 1:** Clinical phenotype of patients tested positive for CASPR2 autoantibodies

| patient number | gender/<br>age at 1<sup>st</sup> contact | clinical phenotype | pain |  | autoanti-bodies | IgG subclass | patient subclassification | serum titer | CSF titer |
| --- | --- | --- | --- | --- | --- | --- | --- | --- | --- |
|  |  |  | widespread | distal burning pain |  |  |  |  |  |
| P1 | 65/m | epileptic seizures, dysautonomia, myoclonus | n/a | n/a | CASPR2 | IgG4 | no pain/IgG4 | 1:10,000 | 1:1,000 |
| P48 | 63/m | epileptic seizures, cognitive deficits, hallucinations | n/a | n/a | CASPR2 | IgG4 | no pain/IgG4 | 1:32,000 | 1:128 |
| P50 | 69/m | cognitive deficits, myoclonus, epileptic seizures | n/a | n/a | CASPR2 | IgG4 | no pain/IgG4 | 1:3,200 | 1:128 |
| P51 | 75/m | singultus | n/a | n/a | CASPR2 | IgG4 | no pain/IgG4 | 1:6,400 | 1:400 |
| P9* | 55/m | epileptic seizures, dystonia | n/a | n/a | CASPR2 | IgG4=IgG3=IgG2 | no pain/IgG4 + IgG1-3 | 1:500,000 | 1:4,000 |
| P34 | 80/m | mnesic deficits, affective lability | n/a | n/a | CASPR2 | IgG4>IgG2 | no pain/IgG4 + IgG1-3 | 1:1,000 | 1:10 |
| P40 | 64/m | epileptic seizures, cognitive deficits | n/a | n/a | CASPR2 | IgG4>IgG2>IgG1 | no pain/IgG4 + IgG1-3 | 1:100,000 | 1:100 |
| P45 | 75/m | mnesic deficits, epileptic seizures | n/a | n/a | CASPR2 | IgG4>IgG2>IgG3 | no pain/IgG4 + IgG1-3 | 1:750,000 | n/a |
| P47 | 72/m | mild cognitive impairment | n/a | n/a | CASPR2 | IgG4>IgG1=IgG2 | no pain/IgG4 + IgG1-3 | 1:3,200 | 1:320 |
| P7 | 61/m | neuromyotonia, pain |  | distal burning pain | CASPR2 | IgG4 | pain/IgG4 | 1:1,000 | 1:10 |
| P19 | 79/m | sleep disorder, depression, gait ataxia |  | distal burning pain | CASPR2 | IgG4 | pain/IgG4 | 1:6,400 | 1:800 |
| P38 | 63/m | myalgia, hallucinations, mnesic deficits | widespread pain: generalized myalgia |  | CASPR2 | IgG4 | pain/IgG4 | 1:100 | n/a |
| P13* | 65/m | myoclonus, mnesic deficits, distal burning pain and back pain |  | burning and tingling pain of the legs, back pain radiating to the legs and feet | CASPR2 | IgG4>IgG2 | pain/IgG4 + IgG1-3 | 1:1,000 | 1:10,000 |
| P32 | 67/m | mnesic deficits, dysautonomia, gait ataxia, epileptic seizures |  | cramps and myalgia of the legs | CASPR2 | IgG4>IgG2=IgG3 | pain/IgG4 + IgG1-3 | 1:1,000 | 1:100 |
| P33 | 45/m | gait ataxia, pain, peripheral neuropathy |  | burning pain of the lower legs and feet | CASPR2 | IgG4>IgG2 | pain/IgG4 + IgG1-3 | 1:1,000 | 1:100 |
| P46 | 72/m | mnesic deficits, neuropathic pain |  | distal burning pain | CASPR2 | IgG4>IgG2=IgG3 | pain/IgG4 + IgG1-3 | 1:3,200 | 1:1,000 |
| P35 | 17/m | pain, radiating ventrally and to the legs |  | pain of the extremities | CASPR2 | IgG1 | pain/IgG1 | 1:320 | n/a |

\*no Ca<sup>2+</sup> imaging performed

### Ethical statement

Experiments using material from patients of the University Hospital Würzburg have been approved by the Ethics Committee of the Medical Faculty, University of Würzburg, Germany (101/20). Patients from external institutes approved the usage of their sera through informed consent within the German Network for Research on Autoimmune Encephalitis and the local ethics committees.

Experiments with animals were approved by the local veterinary authority (Veterinäramt der Stadt Würzburg, Germany) and the Ethics Committee of Animal Experiments, i.e., Regierung von Unterfranken, Würzburg, Germany (licence no. FBVVL 568/200-324/13).

### Cell lines

HEK-293 cells (Human Embryonic Kidney cells; CRL-1573; ATCC–Global Bioresource Center) were used for all *in vitro* experiments. Cells were grown in minimum essential medium (Life Technologies, Darmstadt, Germany) supplemented with 10% fetal bovine serum, L-glutamine (200mM) and 10,000 U/ml penicillin/streptomycin at 37°C and 5% CO_2_.

### Preparation of DRG neurons

Adult DRG neurons were isolated from 12-16 weeks old CD-1 wildtype mice (Charles River, Sulzfeld, Germany), collected in phosphate-buffered saline (PBS), and detached with Liberase TH (5401135001, Roche, Basel, Switzerland) and EDTA for 30min and then with Liberase TM (5401119001, Roche, Basel, Switzerland) and EDTA for 10min at 37°C. After centrifugation (600xg) and trituration, cells were cultured on poly-l-lysine coated coverslips and maintained for two days in Dulbeccos’s Modified Eagle Medium (DMEM)/F12 (1:1) with GlutaMAX™ (31331-028, Gibco, New York, USA) supplemented with 10% fetal calf serum (FCS) and 1% Penicillin/Streptomycin (15140-122, Gibco, New York, USA), at 37°C and 5% CO_2_.

### Microfluidic chambers

Microfluidic chambers (MFCs) (IND150, Xona Microfluidics®, Temecula, USA) were placed on coverslips followed by coating with poly-L-lysine (P2636, Sigma-Aldrich, Burlington, Massachusetts, USA) for 24–48h and laminin-111 (23017-015, Thermo Fisher Scientific, Waltham, Massachusetts, USA) for at least 3h. DRGs were prepared from C57Bl/6 (Jackson Laboratory, Bar Harbor, ME, US) mice at embryonic day 13 (E13) in Hanks’ Balanced Salt Solution (HBSS), and dissociated in trypsin (LS003707, Worthington Biochemical, Lakewood, New Jersey, USA) for 30min at 37°C. Trypsinization was stopped with Neurobasal (NB) Medium (21103-049, Gibco, New York, USA) supplemented with 1% GlutaMAX™ (35050061, Gibco, New York, USA), 2% B27 supplement (17504001, Gibco, New York, USA), and 10% horse serum (16050-122, Gibco, New York, USA). The ganglia were then triturated, and the resulting cell suspension pre-plated for 90min. The cells in the supernatant were then collected and centrifuged at 400×g for 8min. Cell from the pellet were resuspended in 5µl NB medium with 1% GlutaMAX™, 2% B27 supplement and 2% horse serum and seeded in the previously prepared MFC. Cells were kept in NB medium with 1% GlutaMAX™, 2% B27 supplement and 2% horse serum. The somatic compartment contained from day 1 in culture (DIV): 10ng/ml nerve growth factor (NGF) (N-100, Alomone Labs, Jerusalem, Israel), 5ng/ml brain-derived neurotrophic factor (BDNF) (prepared by M. Sendtner, Institute of Clinical Neurobiology, University of Würzburg), 5ng/ml ciliary neurotrophic factor (CNTF) (prepared by M. Sendtner, Institute of Clinical Neurobiology, University of Würzburg) and 5ng/ml recombinant human glial cell line-derived neurotrophic factor (GDNF) (PeproTech, Cranbury, New Jersey, USA). The axonal compartment contained from DIV1: 40ng/ml NGF, 5ng/ml BDNF and 5ng/ml CNTF. To both sides, 1µM 5-Flurodesoxyuridine (FDU) (10124860, Thermo Scientific Chemicals, Waltham, Massachusetts, USA) was added. The medium was exchanged twice by medium without FDU. Patient sera were applied on DIV7 for 2h prior to cell fixation and staining.

### Transfection of cells

HEK-293 cells (CRL-1573; ATCC – Global Bioresource Center, Manassas, VA, USA) were transfected by using a calcium-phosphate precipitation method.^30^ CASPR2 plasmid DNA (1µg/µL; kindly provided by J. Dalmau) was introduced alone or in combination with plasmid DNA (1µg/µL) of either Kv1.1 or Kv1.2 (kindly provided by H.Terlau).

### Immunocytochemistry

For live staining, HEK-293 cells 72h posttransfection or adult DRG neurons at DIV 3 were incubated with commercial anti-CASPR2 antibodies (AF5145, 1:250, R&D Systems, Minneapolis, MN, US) and/or human patient serum (1:50-1:250) in medium for 1h at 4°C. Cells were fixed for 30min at 4°C in 4% PFA/ 4% sucrose in PBS and blocked for 30min at room temperature (RT) with 10% BSA or 5% horse serum in PBS. When needed, cells were also permeabilized with 0.1% Triton-x-100 while blocking. For co-staining of Kv1.1 or Kv1.2 the primary antibodies (ab65790, 1:100, Abcam; 75-008, Cambridge, UK; 1:200, NeuroMab, Davis, CA, US) were diluted in PBS with 5% horse serum and incubated for 1h. The secondary antibodies (713-545-147, 1:500, anti-sheep-Alexa-Fluor-488; 109-165-003, 1:500, anti-human-Cy3; 111-165-003, 1:500, anti-rabbit Cy3; 115-165-003, 1:500, anti-mouse-Cy3, all four from Jackson ImmunoResearch, Ely, UK; ab99772, 1:100, anti-IgG1-FITC, Abcam; 9070-30, 1:100, anti-IgG2-AF488, Southern Biotech, Birmingham, AL, USX; F4641, 1:100, anti-IgG3-FITC, Sigma-Aldrich, Darmstadt, Germany; ab99815, 1:100, anti-IgG4 FITC, Abcam) were incubated in 5% BSA or 5% HS in PBS for 1h at RT. Following three washing steps with PBS, cells were then incubated for 5min at RT with 4’,6-Diamidino-2-phenylindol (DAPI) diluted 1:5000 in PBS. The coverslips were washed in PBS and H_2_O and mounted in Mowiol.

### Confocal microscopy

An Olympus Fluoview ix1000 confocal laser scanning microscope (Olympus, Hamburg, Germany) with an UPLSAPO 60x oil objective (N.A. 1.35), diode lasers of 405nm, 473nm, 559nm and 635nm and the Fluoview FV1000 software was used for imaging. Images were captured in 1024×1024 pixels.

### Super-resolution microscopy

Lattice structured illumination microscopy (SIM^2^) was performed using a Zeiss ELYRA 7 (Carl Zeiss Microscopy GmbH, Jena, Germany) equipped with a Plan-Apochromat 63x/1.40 oil immersion objective and HR Diode 488nm, HR DPSS 561nm and HR Diode 642nm lasers. Z-stacks of images were captured. All images were SIM^2^ processed in the software ZEN 3.0 SR FP2 (Carl Zeiss Microscopy GmbH, Jena, Germany) by a two-step reconstruction algorithm where order combination, denoising and median filtering followed by subsequent iterative deconvolution were performed. Channel alignment via affine transformations generated from z-stacks of embedded TetraspeckTM beads (Z7279, 1:1,000, Thermo Fisher Scientific, Massachusetts, USA) was used for correction of chromatic aberration.

### Nocifensive behavior screen in *Drosophila*

Nocifensive behavior of third instar larvae was analyzed essentially as previously described.^31^ Animals were raised at 29°C and stimulated with a single noxious mechanical stimulus using a 40mN von Frey filament. The immediate behavioral response was classified as positive if the larvae exhibited at least one corkscrew body roll.^32^ Animals with RNAi-mediated knockdown of *neurexin-IV* (*ppk-GAL4*>*UAS-nrx-IV^RNAi^*; BDSC#28715) or *shaker* (*ppk-GAL4*>*UAS-shaker^RNAi^*; VDRC#104474)^33^ specifically in C4da nociceptors were compared to the respective genetic control (*ppk-GAL4/+*). The data were collected from 10 trials (N) each sampling the responses of 30 larvae and analyzed with Prism 9.3 (GraphPad, Prism). An unpaired two-tailed t-test was used to compare the normally distributed data.

### Ca^2+^-imaging

To analyze the changes in spontaneous activity, two days old DRG neurons were subjected for 2h with patient sera or healthy control. For calcium dye loading, DRGs were treated with 5µM Oregon Green BAPTA 1-AM (OGB1, Life Technologies, Carlsbad, CA, USA) for 15min at 37°C. During measurements cells were constantly perfused with HEPES-buffered ACSF (in mM: 4.5 KCl, 2.5 NaH_2_PO_4_, 1 MgCl_2_, 2 CaCl_2_, 120 NaCl, 10 HEPES, 25 glucose, pH 7.4 adjusted with NaOH) using a peristaltic pump at 37°C. A BX51WI upright microscope (Olympus, Hamburg, Germany) equipped with a 20x water-immersion objective UMPLanFL NA 0.5 and a pE-4000 fluorescence illumination system (CoolLED, Andover, U. K.) were used for imaging at 10Hz with a Rolera XR Mono fast 1394 CCD (Qimaging, Surrey, Canada) camera. For each condition 5 videos per culture with 1500 frames (300s) were captured with the software Streampix 4.0 (Norpix, Montreal, Canada) at a rate of 10 frames per second and a binning of 2.

To define regions of interest (ROIs) consisting of DRGs and not the contaminating glial cells, the Fiji plugin “StarDist” was used with a threshold of 0.7 to identify DRGs using star-convex shapes.^34^ The software Bio7 and the Neuron Activity Tool^35^ were used to analyze the fluorescent intensity in AU within the generated ROIs calculating calcium activity peaks/events. Parameters used for analysis were “signal-to-noise” of 2, “average threshold” of 1, “general activity tendency” turned off, “include variance” turned to 30, and “minimum activity counts” of 2. With the number of counted total activities, the spontaneous calcium activity per minute per neuron was calculated.

### Monoclonal CASPR2 autoantibodies

CSF cells from patients with anti-CASPR2 encephalitis were analyzed using single-cell RNA sequencing (scRNA-seq) as previously described.^36^ Single-cell suspensions were prepared and sequenced following standard protocols on Illumina platforms. The variable heavy (VH) and light chain (VL) sequences of clonally expanded B cells were identified using 10x Genomics’ Cell Ranger software, with data referenced to GRCh38. The cloning strategy, vectors, and expression/purification methods were previously described.^36^ Briefly, all sequences were synthesized by GeneArt (Thermo Fisher Scientific, Darmstadt, Germany) and cloned into pcDNA3.1(+) expression vectors. Heavy chain variable regions were subcloned using NheI/PpuMI into an acceptor vector containing an IgG4 heavy constant region. HEK-293 cells were transfected with Lipofectamine 2000 (Life Technologies, Darmstadt, Germany) and 16μg DNA per vector (light and heavy chains). The supernatant was incubated with CaptureSelect™ IgG-CH1 Affinity Matrix (Thermo Fisher Scientific, Darmstadt, Germany), and antibodies were eluted by acidic elution (pH 2.8) and neutralized. Purified recombinant human antibodies (rHumAbs) were desalted using Zeba™ Spin Desalting Columns (Thermo Fisher Scientific, Darmstadt, Germany) and concentrated with a Christ Rotational Vacuum Concentrator System. Antibody concentration was measured using a Qubit protein assay (Q33211, Life Technologies, Darmstadt, Germany). Antigen specificity was confirmed through cell-based assays in HEK-293T cells expression full-length human CASPR2 as previously described.

### Electrophysiological recordings

Patch clamp recordings in a whole-cell configuration from adult DRG neurons at DIV2 were performed at RT to measure the maximal current amplitudes (I_max_) with an EPC-10 HEKA amplifier. Cells were held at -70mV. A voltage step protocol [-80mV, +40mV] with 10mV increments and 2.5s delay was applied. The extracellular solution consisted of [mM]: 135 NaCl, 5 KCl, 2 MgCl_2_, 2 CaCl_2_, 5 glucose, 10 HEPES, pH 7.3 adjusted with NaOH. The intracellular solution consisted of [mM]: 140 KCl, 2 MgCl_2_, 1 CaCl_2_, 2.5 EGTA, 10 HEPES, pH 7.3 adjusted with KOH. Recording pipettes used had a resistance of 3-6MΩ. The current maximum amplitude was measured between 5–35ms of the recorded traces. Cells with a capacitance between 15-35pF were used. Toxins (Conotoxin κM-RIIIJ and α-Dendrotoxin, 100nM) were applied through OctaFlow II system (ALA Scientific Instruments, NY, USA).

### Experimental design and statistical analysis

Short-and long-effects of CASPR2 aAbs were measured through either 2h or 24h incubation with patient sera pools before recording. All experiments were performed in triplicates if not stated otherwise and done blinded for the researcher.

Data from calcium imaging were analyzed using GraphPad Prism, version 10.1.2. Data from electrophysiological recordings were analyzed using RStudio, R version 4.2.2 and GraphPad Prism. SIM^2^ images were analysed by Imaris Software 10.2 using the Spots function, calculating the shortest distance from CASPR2 Spots to Kv1.2 Spots. A threshold of 0.3µm was applied for colocalization. Statistics and plotting were performed using GraphPad Prism. Figures were generated with Origin 10, GraphPad Prism, and Microsoft Office PowerPoint. The thumbnail figure was “Created with BioRender.com”. Data are always represented as mean±S.E.M. (standard error of the mean). Normality of the data was reviewed by Shapiro-Wilk normality test (α=0.05). Statistical significance was calculated using a one- or two-way *ANOVA*. All numbers of experiments (*N*), cells and *p*-values are given in **Supplemental Tables 2-5**. The 0-hypothesis was rejected at a level of *p*<0.05.

### Data availability

Data that support the findings of this study are available from the corresponding author, upon reasonable request.

## Results

### Classification of patients according to the IgG subclass

Our 49 patient samples (40 patient sera were subclassified to their IgG class of CASPR2 aAbs previously^8^) were screened for the predominant IgG subclass and subdivided into four different groups according to their pain phenotype and IgG subclass distribution: patients with pain and IgG4 only, with pain and IgG4+additional IgG1-3 (IgGX), patients without pain and IgG4 only and without pain and IgG4+additional IgG1-3 (IgGX).

CASPR2 aAb positive sera were first confirmed in cell-based assays using CASPR2-transfected HEK-293 cells as well as on adult mouse DRG neurons in costainings with a commercial anti-CASPR2 antibody (**Figure 1A**). Representative stainings of total IgG as well as IgG subclasses of two selected patient sera show a serum positive for IgG4 only (P15) and a serum positive for all IgG subclasses (P23) (**Figure 1B**). Except for three^8^, all patients were IgG4 positive validating IgG4 as the main subclass as previously described.^5,15^ The other three IgG subclasses appeared also quite prominent in the order IgG2>IgG3>IgG1(**Figure 1C**). IgG4 was often detected together with IgG2 (about 65% of all patients) while the presence of IgG4 together with either IgG3 or IgG1 was less prominent (35-37%). The subclass determination allowed for the further classification of our CASPR2-positive samples into four groups including the pain phenotype. Pain and no pain were similarly distributed within the patient cohort (41-51%, **Figure 1D**). In 7% of patients, the pain status was not recorded. Pain and no pain patients were subclassified into IgG4 as the only subclass (further on labeled IgG4) present and IgG4 with at least one additional IgG which can be IgG1 and/or IgG2 and/or IgG3 (IgGX). Pain and no pain patients with IgG4 only were less representative (7 and 17%) than patients with CASPR2 aAbs with IgG4 and at least one additional IgG (34%; **Figure 1D**). Seventeen selected patient serum samples were used in the present study to further study the CASPR2 aAb pathophysiology (**Table 1**).

**Figure 1.**
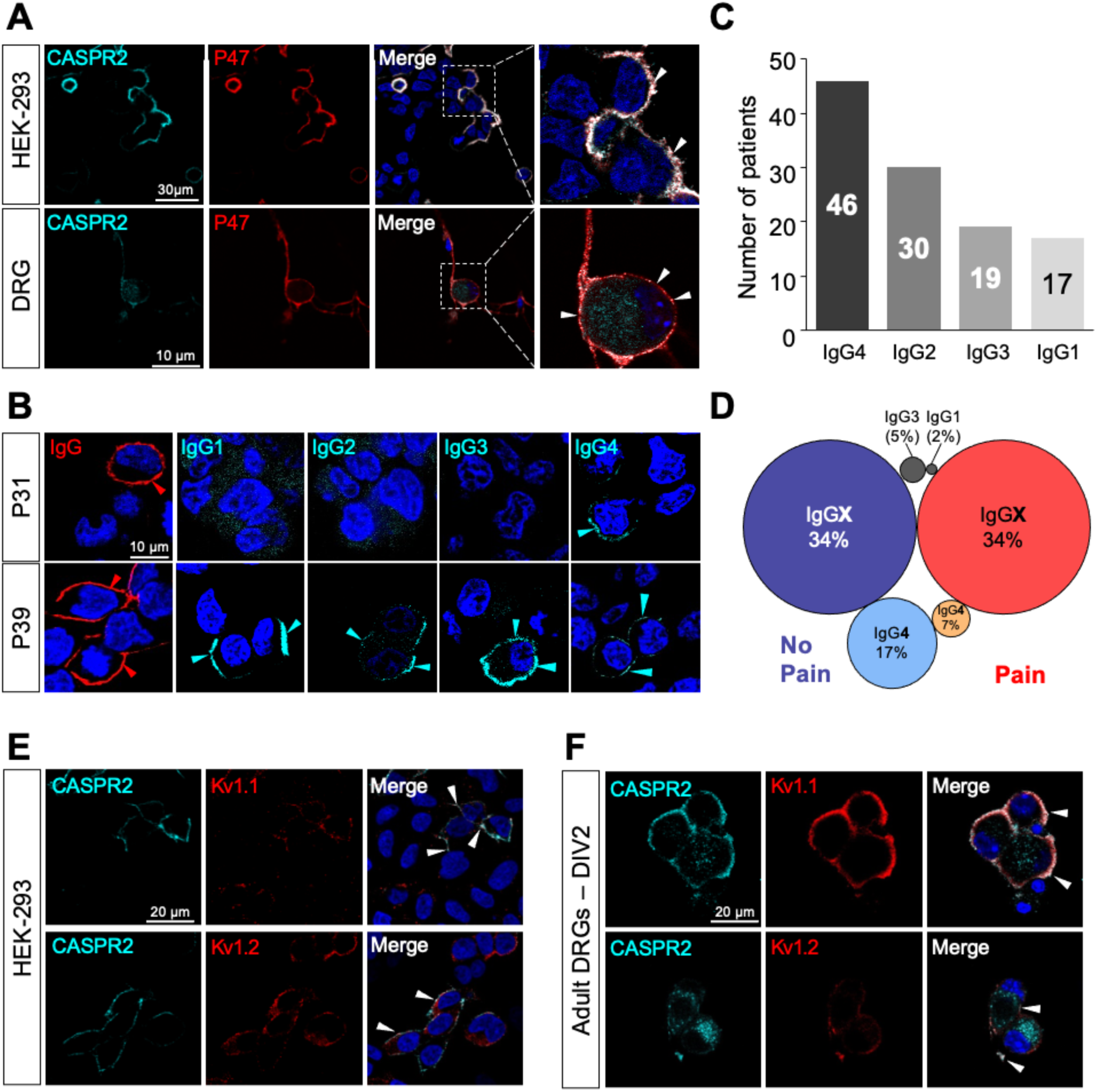
CASPR2 aAb in patient serum show different IgG subclass compositions. A) Exemplary immunocytochemical stainings of CASPR2 (cyan) and patient serum (red) binding to membrane of CASPR2 transfected HEK-293 cells and adult DRG neurons. B) Stainings for different IgG subclasses (cyan) in two exemplary total IgG positive (red) patient sera. C) Number of patients positive for different IgG subclasses (*n*=49). D) Distribution of IgG subclasses (IgG4 only or IgG4 plus additional IgG = IgGX) and pain phenotype (with pain - red and orange circles, no pain – light and dark blue circles with circle size related to number of positive patient sera. E, F) Costainings of CASPR2 (cyan) with either Kv1.1 (red) or Kv1.2 (red) in cotransfected HEK-293 cells or adult DRG neurons. Colocalization of CASPR with Kv channels are marked by white arrow heads (merged images).

### CASPR2 expression is unaltered in DRGs upon presence of anti-CASPR2 aAbs

CASPR2 belongs to the VGKC expressed at juxtaparanodal regions at the node of Ranvier in the PNS. It is also functionally expressed in DRG neurons, which represent active mediators in the development of neuropathic pain to transmit pain signals from the PNS to the CNS.^23,24^ Within the VGKC, CASPR2 interacts with potassium channels of the Kv subtype via proteins 4.1b and PSD95. To further study the protein complex formation in the presence of CASPR2 aAbs, colocalization of CASPR2 with Kv1.1 or Kv1.2 was verified in transfected HEK-293 cells and adult and embryonic (E13) DRG neurons at DIV2 (**Figure 1E, F, Supplemental Figure 1**).

The pathophysiology of other aAbs includes changes in the expression pattern of the targeted protein.^37,38^ CASPR2 has been determined with a protein half-life in transfected HEK-293 cells of 3.7h.^39^ Patients harbor CASPR2 aAbs, however, for days or even weeks before a clinical phenotype manifests. Moreover, CASPR2 aAbs are mainly of the IgG4 subclass unable to crosslink proteins and induce their subsequent internalization. To mimic disease, CASPR2 expression was analyzed in DRG neurons incubated for one day (1d), 2d, and 4d with patient sera. To model plasma exchange, we exchanged the serum after 2d with a healthy control serum for 2d (2R for rescue group) (**Figure 2A, Supplemental Figure 2A**). CASPR2 expression was estimated as relative CASPR2 density per 100µm of axon length using ten patient sera and a healthy control serum individually. Group analysis according to the pain phenotype independent of which IgG subclass revealed a significant decrease in CASPR2 expression between day 1 and day 4 for the no pain group (*p*=0.0147) but not in the pain group (*p*=0.9536) or for different IgG subclasses (IgG4 *p*=0.1413; IgGX *p*=0.4255; **Figure 2B, Supplemental Table 1**).

**Figure 2.**
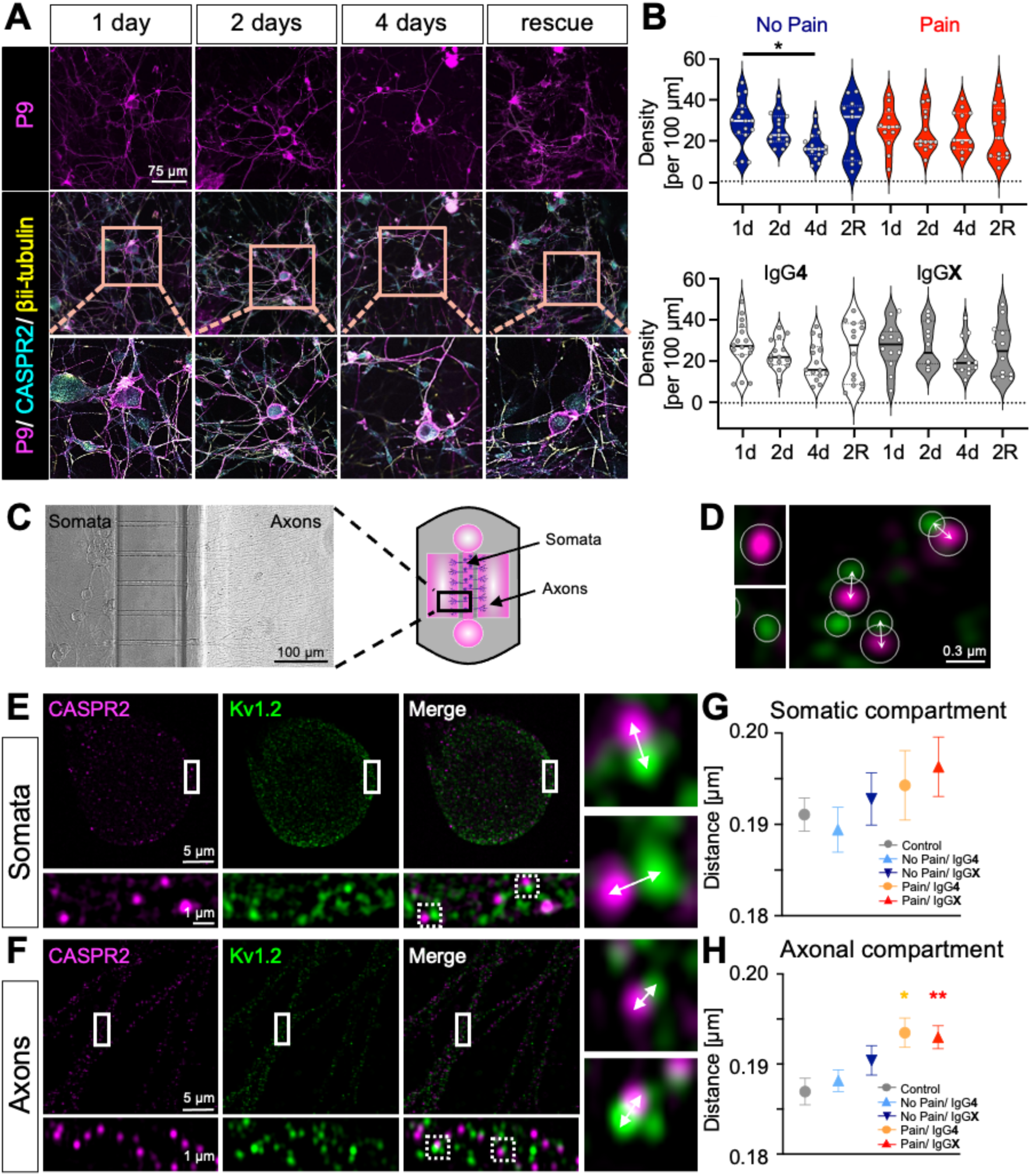
Expression of CASPR2 after exposure to CASPR2 aAbs and structural integrity of the VGKC. A) Experimental design of CASPR2 aAbs incubation on DRG neurons for 1d, 2d, 4d, or rescue with 2d aAb exposure and 2d withdrawal (2R). A) Immunocytochemical staining of DRG neurons for patient serum (magenta), CASPR2 (cyan) and bIII-tubulin (yellow). B) Quantification of relative CASPR2 expression density per 100µm axon for four patient serum pools with (red) or without pain (blue) and with only IgG4 (white) or additional other IgG subclasses (IgGX, grey) after different time points. Data are shown as violin plots with individual values, median=bold line, quartiles=dotted lines. *N*=4-5, *n*=12-16. C) Microfluidic chamber with embryonic DRGs to separate somatic and axonal compartments and investigate VGKC organization after CASPR2 aAbs incubation in separated compartments. D) Exemplary pictures of image analysis showing non-colocalizing CASPR2 (magenta; top left) and Kv1.2 (green; bottom left) signals, and colocalizing (right). Arrows indicate distance measurement with a threshold of 0.3µm. E, F) Exemplary pictures using lattice SIM^2^ microscopy of soma (E) and axons (F) from DRGs in microfluidic chambers treated with healthy control serum. Arrows mark the distance between CASPR2 (magenta) and Kv1.2 (green). G, H) Quantification of distances between CASPR2 and Kv1.2 for somata (G) and axons (H) after CASPR2 aAbs incubation for 2h. Data are shown as mean±SEM. Somatic and axonal complexes: *n*=325-1465 and *n*=1872-3603 respectively. Levels of significance: \**p*<0.05, \*\**p*<0.01.

An analysis of CASPR2 density considering patient serum samples from both no pain groups (IgG4 and IgGX) separately exhibited a lower CASPR2 density on days 2 and 4 compared to day 1. In contrast, the pain group with IgG4 revealed a slight increase in CASPR2 density along the axons (**Supplemental Figure 2B**). However, all four groups individually analyzed did not significantly change CASPR2 protein abundance (**Supplemental Figure 2B**). In sum, the CASPR2 expression alterations in DRG neurons supplemented with CASPR2 aAbs were minor and most likely did not explain a causative correlation to the IgG subclass present.

In a comparative approach, we used an *in vivo* setting to quantify nocifensive behavior elicited by mechanical *von Frey* filament stimulation in *Drosophila* larvae^31,32^. Whereas knockdown of the voltage-gated potassium channel *shaker* (*Drosophila* Kv1 homolog) via nociceptor-specific RNAi gave a phenotype resembling hyperalgesia, *neurexin-IV* (*nrx-IV*) knockdown in fact slightly decreased nocifensive behavior (**Supplemental Figure 3**). These data emphasize the evolutionarily conserved role of voltage-gated potassium channels in nociceptors and indicate that the potassium channel subunits of the VGKC represent excellent candidates for the nociceptive association of CASPR2 aAbs.

### The structural organization of the VGKC is altered in the presence of anti-CASPR2 aAbs from patients with pain

Possible structural re-arrangements of CASPR2 and Kv potassium channel subunit proteins of the VGKC were investigated with high-resolution SIM^2^ imaging. MFCs have been used to grow DRG neurons (**Figure 2C**) to improve resolution by clear discrimination between somatic and axonal localization. Somatic and axonal compartments were both incubated separately with four different patient serum pools depending on pain phenotype and IgG subclass for 2h (**Table 1**). Afterwards, CASPR2 and Kv1.2, both belong to the VGKC, were stained and analyzed for their distances (**Figure 2D-H**). While no significant distance changes were obtained for the somatic compartment, a significant increase in the distances between CASPR2 and Kv1.2 was seen for both pain groups independent on the IgG composition compared to healthy control (**Figure 2F, H, Supplemental Table 2**). In sum, we only observed structural changes in the VGKC complex upon incubation with CASPR2 aAbs from patients with pain.

### Assessment of neuronal activity after incubation with anti-CASPR2 by Ca^2+^-imaging

To control for functional alterations and neuronal activity, the excitability of adult DRG neurons was investigated in Ca^2+^-imaging experiments. Two hours after CASPR2 aAb treatment, we analyzed the spontaneous activity of adult DRG neurons (**Table 1, Supplemental Table 3**). The fluorescent intensity traces from representative cells from all five conditions and additional heatmaps of ∼40 cells are shown (**Figure 3A**). After 2h of incubation, again only the two groups associated with pain harboring either IgG4 or IgGX CASPR2 aAbs significantly increased the neuronal activity of DRG neurons measured by increasing calcium transient frequency, amplitude and area under curve (AUC) when compared to incubation with healthy control serum (frequency: pain/ IgG4 *p*<0.0001; pain/ IgGX *p*=0.0100; amplitude: pain/ IgG4 *p*=0.0343; pain/ IgGX *p*=0.0391; AUC: pain/ IgG4 *p*=0.0180; **Figure 3B, Supplemental Table 3**). Taken together, the hyperexcitability observed for adult DRG neurons after treatment with CASPR2 aAbs from patients suffering from neuropathic pain points towards a significant impact on impaired spontaneous activity at the DRG neurons.

**Figure 3.**
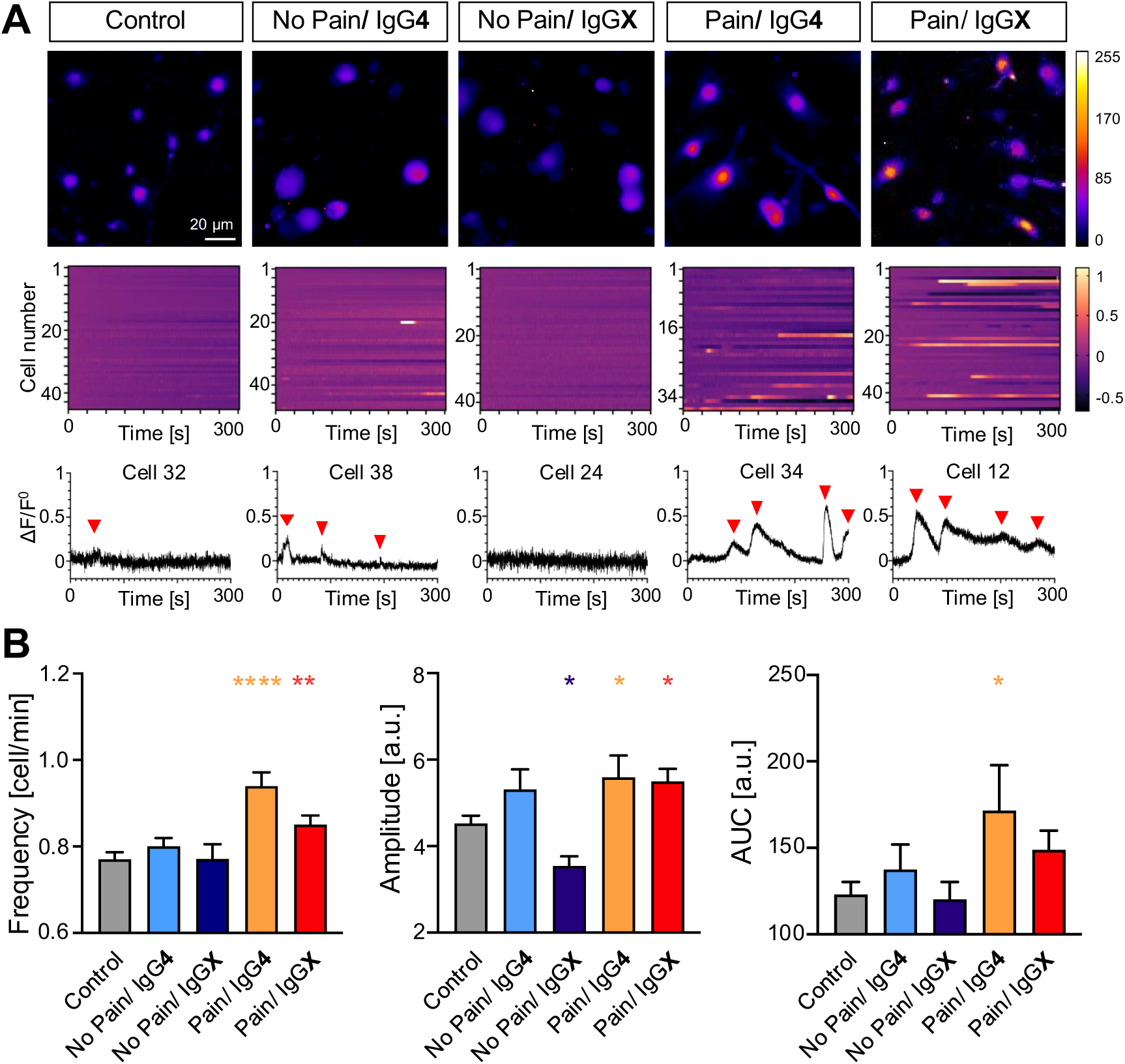
Spontaneous calcium activity of DRGs after incubation with CASPR2 aAbs. A) Exemplary pictures and activity graphs of ROIs after 2h incubation with control serum or patient serum. Neuronal activity from ∼40 cells is shown in a heatmap. B) Frequency (left), amplitude (middle) and area under curve (AUC; right) of spontaneous calcium transient activity events after incubation with CASPR2 aAbs of different serum subclassifications. Data shown as mean ± SEM, *n*=718 (control), *n*=682 (no pain/IgG4), *n*=476 (no pain/IgGX), *n*=566 (pain/IgG4), *n*=727 (pain/IgGX). Levels of significance: \**p*<0.05, \*\**p*<0.01, \*\*\*\**p*<0.0001.

### Effects of anti-CASPR2 aAbs on the function of the associated Kv channels

CASPR2 aAbs are known to decrease the function of the associated Kv channel within the VGKC leading to hyperexcitability of DRG neurons.^13^ However, the sample size was small (n=2 CASPR2-positive sera) and IgG subtypes were not reported. Therefore, we now investigated the function of the associated Kv channels in medium-sized adult DRG neurons after a 2h exposure to CASPR2 aAbs.

All four subgroups of CASPR2 positive patient serum samples significantly decreased the function of Kv channels (for voltage range -50 to -10 *p* <0.05 for all groups except pain/IgGX, from 0 to 40 IgG4 groups *p*<0.01, **Figure 4A**). The most prominent impairment of potassium currents was observed for the groups with only IgG4 independent of the pain phenotype after 2h (**Figures 4B-E**). Decreased Kv channel activity after short-term incubation was noticed for all four groups of patients even if carrying IgG4 and an additional IgG, suggesting IgG4 as the main driver of hyperexcitable DRG neurons in the presence of CASPR2 aAb-containing sera. We next selected one patient with non-IgG4 CASPR2 aAbs and did not observe alterations of the potassium channel activity after 2h pre-treatment with patient serum, further providing strong evidence for IgG4 as the key IgG of CASPR2 autoantibody pathophysiology (**Figure 4F**). To verify a causal effect of aAb binding, the autoantibodies were withdrawn by serum exchange for 24h following 2h presence of the CASPR2 aAbs. This resulted in a return to baseline condition. Incubation with CASPR2 aAbs for 24h also caused a reduction of the Kv channel activity although less prominent (**Supplemental Figure 4**). Hence, our data reveal a local and direct effect of CASPR2 autoantibodies on the excitability of the sensory neurons mediated by impairing Kv channel activity (**Figure 4G, H, Supplemental Table 4**). The recorded traces at different voltage steps (−80mV to +40mV) had different current patterns. This may point towards a contribution of Kv1.1, Kv1.2, and possibly Kv1.1/Kv1.2 heteromers but also argues for other Kv subtypes present. Using the specific toxin κM-RIIIJ which specifically blocks Kv1.2 heteromers, we could confirm that Kv1.2 heteromers represent a significant portion of the recorded ion channels (*p*<0.001). A similar portion of potassium currents could be blocked with α-dendrotoxin which blocks Kv1.1, Kv1.2, Kv1.6 (*p*<0.001, **Figure 4I, J, K, Supplemental Table 5**). To confirm, that the effect was indeed mediated specifically by CASPR2-binding aAbs and not by aAbs targeting other neuronal components present in polyclonal patient sera, we utilized monoclonal anti-CASPR2 aAbs isolated from antibody-secreting cells in patients’ CSF cloned to an IgG4 backbone. The IgG4 backbone was modified to prevent Fab arm exchange and thus restore a cross-linking ability to CASPR2 IgG4 aAbs^40,41^. The monoclonal antibodies used were known to target either the discoidin domain which has been described as a main target of CASPR2 aAbs or the laminin domain which is present four times in the extracellular domain of CASPR2 (**Figure 5A**). Both monoclonal anti-CASPR2 antibodies indeed reduced the potassium channel activity but the effects were less prominent than the patient serum pools of the subclassified patients. The decreased potassium channel current was mainly observed at negative potentials between -80 to -40 mV (**Figure 5B, C**). An impairment of potassium channel function has also been demonstrated for aAbs against another adhesion protein LG1 (against LRR domain)^42^ which was used as a positive control (**Supplemental Figure 5**).

**Figure 4.**
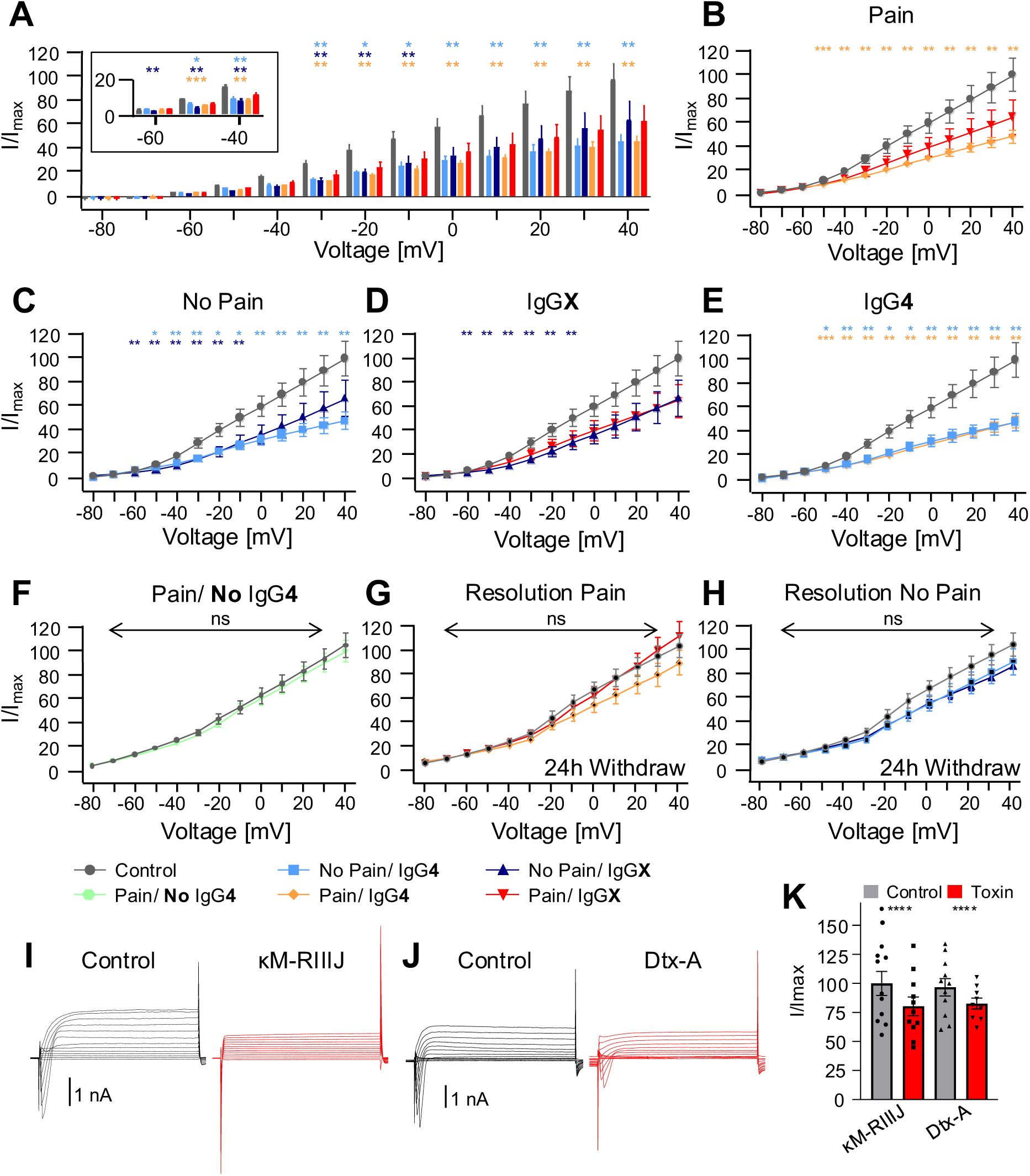
Decrease of Kv currents upon CASPR2 aAbs application. A) Bar plot of potassium channel activity at different voltage steps from -80 to +40 mV. B) I-V (current-voltage relation) plot measured after 2h aAb presence on DRG neurons under pain condition; C) no pain condition; D) IgGX, E) IgG4 with *N*=3-4, *n*=9-14, and F) pain/no IgG4, *N*=2, *n*=10. G-H) Withdrawal of the CASPR2 aAbs led to a baseline shift of the potassium channel current (patients with pain and patients without pain), *N*=2, *n*=10-11. I-K) Representative potassium channel current traces as well as the effect of specific potassium channel subunit blocker (α-dendrotoxin Dtx-A and κM-RIIIJ (red traces) shown in the bar plot (at +40 mV). Data shown as mean±SEM. Levels of significance: \**p*<0.05, \*\**p*<0.01, \*\*\**p*<0.001, \*\*\*\**p*<0.0001.

**Figure 5.**
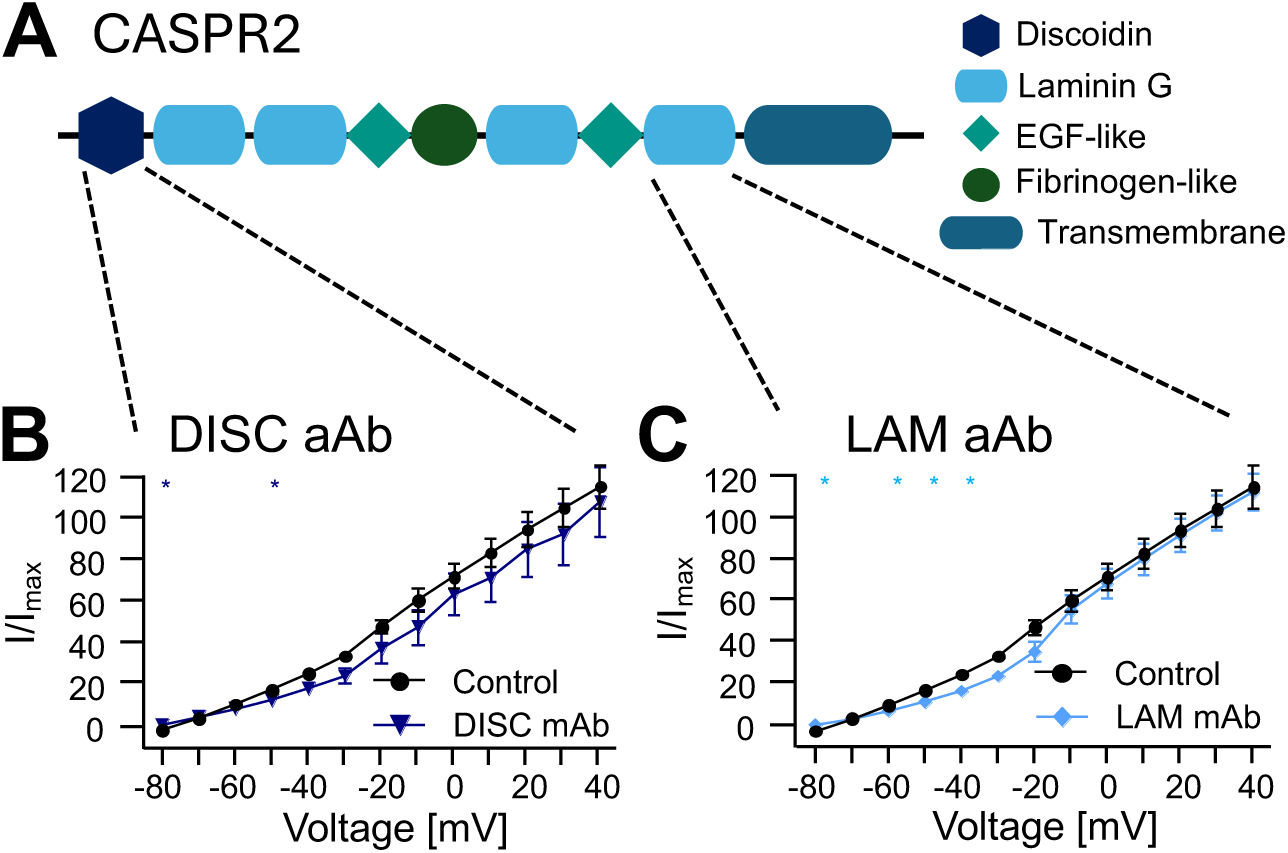
CASPR2 monoclonal autoantibodies isolated from patients also decrease potassium channel activity in DRG neurons. A) Domain architecture of CASPR2 including the Disc domain, fibrinogen-like domain, EGF-like domains, and laminin-domains. B) Patch clamp recordings in the presence of patient-derived monoclonal CASPR2 autoantibodies, one against the Disc and the other against the laminin domain using a voltage step protocol from -80 to +40 mV demonstrates a significant alteration of the potassium channel activity especially at negative potentials. *N*=2, *n*=11. Level of significance: \**p*<0.05.

In summary, we document a direct effect of CASPR2 aAbs of the IgG4 subtype on the function of the associated Kv channel. IgG4 is the key driver of increased excitability of sensory DRG neurons, a pathophysiological effect more pronounced in the presence of a pain phenotype associated with CASPR2 aAbs from patient sera.

## Discussion

CASPR2 aAbs have been found in patients with various neurological diseases including neuropathic pain. We subclassified patient sera positive for CASPR2 aAbs according to their pain phenotype and IgG subclass which enabled us to shed further light on differences in pathological mechanisms of those aAbs and their association with pain.

CASPR2 is an adhesion protein and part of the VGKC. A previous study documented an increased pain sensitivity following passive transfer of purified patient CASPR2 IgG from two patients into mice.^13^ Patient-derived CASPR2 IgG increased DRG excitability but only in one patient while aAbs from the other patient lacked this effect. Decreased CASPR2 and Kv levels were detected at the juxtaparanodal regions from injected animals.^13^ In contrast, others have described an enhanced cluster formation of CASPR2, an increased expression of the associated potassium channel subunit Kv1.2, but an almost unaffected CASPR2 expression in transfected HEK-293 cells and cultured hippocampal neurons.^11^

CASPR2 aAbs have been determined as mainly of the IgG4 subclass^5,14,15^, hence differing from the other IgG classes by their inability to activate complement, crosslink proteins and induce their subsequent internalization. Recently, we found that the majority of patients with CASPR2 aAbs harbor aAbs not only of the IgG4 subclass but at least one other IgG subclass in addition.^8^ Thus, the additional subclass might contribute to differences in the pathological mechanisms.

Besides the importance of the IgG subclass, even domain-specific targeting effects of aAbs may play a role. Such domain-specific pathophysiological changes have been described from patient-derived aAbs against the adhesion protein LG1 and changes in the spatial organization of associated potassium channels.^42^

Simulating the situation in the patient before and after (plasmapheresis) treatment using a time course of four days and a subsequent recovery period of two days after aAbs withdrawal, we found a slight decrease in endogenous CASPR2 expression in treated DRG neurons for patient subclassifications (pain versus no pain and IgG4 versus IgGX). The observed decrease in CASPR2 expression after four days of presence of aAbs was however only significant in the no pain groups independent of the IgG subclass. These data align with unaltered or minor decrease in CASPR2 expression in primary neurons or following passive transfer of patient IgG in mice.^11,13^ After an exchange of the CASPR2 aAbs to a healthy control serum, expression levels were clearly rescued to baseline level independent of patient subclassifications. The half-life of CASPR2 has been determined to be 3.7h^39^, arguing that within several magnitudes of protein turnover, this slight decrease in CASPR2 expression is unlikely to be the main pathological function of CASPR2 aAbs.

Further structural analysis using high-resolution SIM^2^ microscopy revealed a significant increase in the distances between CASPR2 and Kv1.2, both being members of the VGKC, along DRG axons but not at the soma in the presence of CASPR2 aAbs from patients with pain suggesting that the protein complex loses its structural integrity upon aAb binding. The VGKC structural integrity was however unchanged for patients without pain. The non-persisting structural integrity of the VGKC upon CASPR2 aAb binding is thus similar to other aAb against adhesion proteins e.g. LG1, where a spatial reorganization of the associated Kv channels along the axon initial segment has been found to underlie functional impairment of the affected neurons.^42^ We further concentrated on the mechanism of pain association with CASPR2 aAbs which has been suggested to occur via hyperexcitability of DRG neurons^13^ as mediators of pain signaling to the CNS. In *Drosophila* larvae, a RNAi-mediated knockdown of the CASPR2 homolog *nrx-IV* did not elevate nocifensive behavior. In contrast, knockdown of the potassium channel subunit *shaker*, the mammalian Kv1 homolog, led to an increase in nocifensive responses. This is in line with potassium channels controlling neuronal excitability^43,44^ being involved in mechanisms underlying neuropathic pain in patients with CASPR2 aAbs.

Using Ca^2+^-imaging in order to study neuronal excitability, only patient serum samples associated with a pain phenotype significantly increased the frequency, the amplitude and area under the curve of measured calcium transients. The neuronal activity pattern was almost unchanged for serum samples without pain association. If increased neuronal activity is a direct consequence of exposure to pain-associated CASPR2 aAbs and thus the functionality of the VGKC, similar effects would be expected from patch clamp recordings considering Kv channel conductance. Interestingly, whole-cell recordings from pre-treated DRG neurons with CASPR2 aAbs from patient subclassifications showed significantly decreased potassium channel activity for all patient serum subclasses with the most prominent effect of the no pain/pain IgG4 subclasses. Depletion of aAbs always rescued potassium channel activity to baseline level indicating that functional alterations of recorded DRG neurons were a direct consequence of aAb binding to the VGKC which is present in almost all DRG subtypes.^26^ Moreover, the presence of an additional IgG class slightly minimized functional DRG impairments. Similarly, CASPR2 aAbs of the IgG4 group had a more pronounced effect on the overall neuronal activity in Ca^2+^-imaging strongly indicating that IgG4 is the main driver of DRG hyperexcitability. A decreased potassium channel activity has been previously shown but could only been observed for one patient sample^13^ proposing that both patients may have been differed in the IgG compositions of CASPR2 aAbs. Elucidating which Kv subtypes may underlie the observed decreased potassium channel function revealed a partial contribution of Kv1.1, Kv1.2 and Kv1.6, which are expressed in virtually all types of DRG neurons.^26,28,45^ Considering epitope-specific pathophysiological effects of aAbs similar to recent findings for LG1 aAbs involving a spatial re-distribution of Kv channels and concomitant impaired neuronal control of action potential initiation and synaptic integration^42^, epitope-specific effects of discoidin and laminin domain targeting patient-derived CASPR2 aAbs on Kv dysfunction were not observed.

Our argument that IgG4 is the key driver of decreased potassium channel activity was further reinforced when using a patient serum lacking IgG4 and exhibiting no alteration of potassium channel function. These data indicate that the pathophysiology of CASPR2 IgG4 possibly by its inability to crosslink two CASPR2 proteins but binding to one CASPR2 protein of the VGKC structurally hinders the activation of the associated potassium channels without affecting the overall structure of the protein complex as such. However, the pathophysiology of neuropathic pain in patients with CASPR2 aAbs includes additional molecular pathways. Disorders due to neuronal hyperexcitability with and without pain have been prone to display an interplay between impaired sodium channel and potassium channel activity leading to decreased thresholds of action potential firing or changes such as prolonged repolarization phases of action potentials.^46,47^ Taken together, the subclassification of CASPR2-positive patient serum samples resembles a key strategy to unravel discrepancies in the pathological mechanisms that are the prerequisite for targeted treatment.

### Conclusions

In sum, our CASPR2 aAb subclassification of patients identified differences in the pathophysiology between the presence of different CASPR2 IgGs with IgG4 as the key immunoglobulin for subsequent neuronal hyperexcitability of sensory DRG neurons. Moreover, the pathophysiology of CASPR2-positive patients is associated with pain and this correlates significantly with a decrease of the structural integrity of the targeted protein complex and consequently an increase of the overall neuronal excitability of cultured sensory neurons. Additional pathomechanisms underlying this increased neuronal activity in pain patients may include functional alterations of other ion channels or specific neuronal subtypes, inflammatory pathways, or the role of immune complexes triggering secondary intracellular signal cascades further manifesting with an associated pain phenotype.

## Abbreviations

(CASPR2): Contactin-associated protein 2
(IgG): immunoglobulin G
(Kv channel): voltage-gated potassium channel
(VGKC): voltage-gated potassium channel complex
(DRG): dorsal root ganglia

## Acknowledgements

We would like to thank Christine Schmitt and Dana Wegmann for excellent technical assistance and Ricarda Hesse for help with the *Drosophila* experiments. Dr. Shrinivasan Raghumaran (University of Utah, Salt Lake City, UT, US) is highly acknowledged for providing the conotoxin to identify potassium channel compositions. We further thank Dr. Heinz Terlau, University of Kiel, Germany and Dr. Josep Dalmau, SJD Barcelona Children’s Hospital and Hospital Clinic, Spain for the expression plasmids of the potassium channel subunits and CASPR2. We are especially thankful to the GENERATE network supporting our project. Fly stocks obtained from the Bloomington *Drosophila* Stock Center (NIH P40OD018537) and the Vienna *Drosophila* Resource Center (VDRC, www.vdrc.at) were used in this study.

## Author contributions

Participated in research design – Carmen Villmann, Kathrin Doppler, Claudia Sommer, Heike Rittner, Michael Briese, Michael Sendtner. Conducted experiments – Margarita Habib, Patrik Greguletz, Anna-Lena Wiessler, Michéle Niesner, Maximilian Koch, Zare Abdelhossein, Mareike Selcho, Ligia Abrante, Christian Werner, Frank Leypoldt, Robert Handreka, Harald Prüss, Jan Lewerenz, Justina Dargvainiene, Péter Körtvélyessy, Dirk Reinhold, Franziska Thaler, Kalliopi Pitarokoili - Performed data analysis – Margarita Habib, Patrik Greguletz, Anna-Lena Wiessler, Mareike Selcho, Robert J. Kittel, Robert Blum, Annemarie Sodmann, Christian Werner. Wrote the manuscript – Carmen Villmann, Kathrin Doppler with the help from coauthors. Revised the manuscript –Kathrin Doppler, and Carmen Villmann.

