## Supplemental information for "Neuropathic pain and distinct CASPR2 autoantibody IgG subclasses drive neuronal hyperexcitability"

### SUPPLEMENTAL TABLES

**Supplemental Table 1: Relative CASPR2 expression density per 100 µm axon after long-term exposure to CASPR2 autoantibodies**

| | single subgroup analysis | | | |
|---|---|---|---|---|
| | Pain IgG4 | Pain IgGX | No Pain IgG4 | No Pain IgGX |
| 1d | 30.5±3.0 | 24.5±4.1 | 30.7±3.17 | 33.3±3.65 |
| 2d | 21.5±4.3 | 27.0±3.6 | 24.3±1.85 | 30.24±5.96 |
| 4d | 28.0±4.4 | 23.2±3.3 | 16.02±2.09 | 18.4±1.33 |
| 2R | 24.8±7.95 | 24.9±4.0 | 24.3±1.85 | 31.5±6.7 |
| *n* | 4 | 8-9 | 10-12 | 4 |
| *N* | 1 | 3 | 3 | 1 |
| | subgroup analysis according to Pain and IgG subclass | | | |
| | No Pain | Pain | IgG**4** | IgG**X** |
| 1d | 31.4±2.5 | 26.56±3.1 | 30.4±2.3 | 27.3±2.96 |
| 2d | 25.5±2.07 | 25.4±2.79 | 23.9±1.72 | 27.58±2.96 |
| 4d | 16.8±1.52 | 24.7±2.67 | 21.3±2.51 | 21.5±2.3 |
| 2R | 26.0±3.94 | 24.8±3.64 | 24.66±3.62 | 28.8±3.86 |
| *n* | 13-16 | 12-13 | 15-16 | 12-13 |
| *N* | 4 | 4 | 5 | 5 |

*N*=number of experiments; *n* =number of analyzed axons

**Supplementary Table 2: Distance calculations between fluorescent signals of CASPR2 and Kv1.2 in the VGKC**

| Compart-ment | Group | Significance | Adj. *p*-Value | Mean | SEM | *n* |
|---|---|---|---|---|---|---|
| Somaic | Control | | | 0.1911 | | 1465 |
| | No Pain/ IgG4 | ns | 0.9694 | 0.1894 | 0.003118 | 754 |
| | No Pain/ IgGX | ns | 0.9724 | 0.1928 | 0.003333 | 620 |
| | Pain/ IgG4 | ns | 0.8991 | 0.1943 | 0.004265 | 325 |
| | Pain/ IgGX | ns | 0.4623 | 0.1963 | 0.003658 | 480 |
| Axonal | Control | | | 0.1869 | | 2327 |
| | No Pain/ IgG4 | ns | 0.9178 | 0.1882 | 0.001895 | 3603 |
| | No Pain/ IgGX | ns | 0.3256 | 0.1904 | 0.002213 | 1872 |
| | Pain/ IgG4 | * | 0.0102 | 0.1935 | 0.002187 | 1952 |
| | Pain/ IgGX | ** | 0.0075 | 0.193 | 0.001957 | 3085 |

ns=non-significant, ; *p<0.05, **p<0.01, *n* =number of colocalized CASPR2 signals

**Supplemental Table 3: Calcium imaging data analysis**

| | Group | Significance | *p*-Value | Mean | SE of diff. | *n* |
|---|---|---|---|---|---|---|
| **Frequency** | Control | | | 0.7708 | | 718 |
| | No Pain/ IgG4 | ns | 0.3402 | 0.8009 | 0.03158 | 682 |
| | No Pain/ IgGX | ns | 0.9845 | 0.7714 | 0.03491 | 476 |
| | Pain/ IgG4 | **** | <0.0001 | 0.9399 | 0.0332 | 566 |
| | Pain/ IgGX | ** | 0.01 | 0.8509 | 0.03108 | 727 |
| **Amplitude** | Control | | | 4.527 | | 718 |
| | No Pain/ IgG4 | ns | 0.1037 | 5.307 | 0.4787 | 682 |
| | No Pain/ IgGX | * | 0.0492 | 3.486 | 0.5292 | 476 |
| | Pain/ IgG4 | * | 0.0343 | 5.593 | 0.5033 | 566 |
| | Pain/ IgGX | * | 0.0391 | 5.5 | 0.4711 | 727 |
| **AUC** | Control | | | 122.9 | | 718 |
| | No Pain/ IgG4 | ns | 0.4579 | 137.4 | 19.52 | 682 |
| | No Pain/ IgGX | ns | 0.0601 | 82.7 | 20.52 | 476 |
| | Pain/ IgG4 | * | 0.0018 | 171.5 | 21.58 | 566 |
| | Pain/ IgGX | ns | 0.179 | 148.8 | 19.21 | 727 |

AUC=area under curve, ns=Non-significant; *p<0.05, **p<0.01, ****p<0.0001; *n*=number of cells analyzed

**Supplementary Table 4: Electrophysiological properties of potassium currents activated from DRG neurons upon CASPR2 autoantibody presence**

| aAb | Voltage | Significance | *p*-Value | Mean 1 | Mean 2 | SE of diff. | *N* (C; sub) | *n* (C; sub) | Res. | Voltage | Significance | *p*-Value | Mean Ctrl | Mean aAb | SE of diff. | *N* (C; sub) | *n* (C; sub) |
|---|---|---|---|---|---|---|---|---|---|---|---|---|---|---|---|---|---|
| Control vs. No Pain/ IgG4 | -80 | ns | 0.2363 | -0.1188 | -0.1596 | 0.03347 | | | Control vs. No Pain/ IgG4 | -80 | ns | 0.2697 | -0.1073 | -0.07671 | 0.02693 | | |
| | -70 | ns | 0.3179 | 0.01248 | 0.009866 | 0.002555 | | | | -70 | ns | 0.5865 | -0.002649 | 0.005295 | 0.01435 | | |
| | -60 | ns | 0.6554 | 0.2311 | 0.2137 | 0.03821 | | | | -60 | ns | 0.3172 | 0.1268 | 0.09776 | 0.02822 | | |
| | -50 | * | 0.0457 | 0.6187 | 0.4195 | 0.09378 | | | | -50 | ns | 0.1907 | 0.2727 | 0.2019 | 0.05208 | | |
| | -40 | ** | 0.0098 | 1.147 | 0.6513 | 0.1735 | | | | -40 | ns | 0.1345 | 0.4469 | 0.3169 | 0.08266 | | |
| | -30 | ** | 0.0072 | 1.91 | 0.9739 | 0.3102 | | | | -30 | ns | 0.1202 | 0.6508 | 0.461 | 0.1155 | | |
| | -20 | * | 0.0101 | 2.7 | 1.371 | 0.4653 | 4; 3 | 14; 9 | | -20 | ns | 0.2546 | 1.049 | 0.8146 | 0.1991 | 2; 2 | 11; 10 |
| | -10 | * | 0.0112 | 3.426 | 1.765 | 0.5922 | | | | -10 | ns | 0.1754 | 1.45 | 1.106 | 0.2442 | | |
| | 0 | ** | 0.0087 | 4.11 | 2.079 | 0.6961 | | | | 0 | ns | 0.1893 | 1.759 | 1.37 | 0.2854 | | |
| | 10 | ** | 0.0067 | 4.826 | 2.383 | 0.804 | | | | 10 | ns | 0.2204 | 2.051 | 1.638 | 0.3259 | | |
| | 20 | ** | 0.0056 | 5.547 | 2.671 | 0.9198 | | | | 20 | ns | 0.2562 | 2.334 | 1.908 | 0.3638 | | |
| | 30 | ** | 0.0047 | 6.275 | 2.964 | 1.034 | | | | 30 | ns | 0.2986 | 2.608 | 2.18 | 0.4005 | | |
| | 40 | ** | 0.004 | 7.01 | 3.237 | 1.147 | | | | 40 | ns | 0.3407 | 2.88 | 2.448 | 0.4414 | | |
| Control vs. No Pain/ IgGX | -80 | ns | 0.1359 | -0.1188 | -0.07627 | 0.02729 | | | Control vs. No Pain/ IgGX | -80 | ns | 0.3267 | -0.1073 | -0.07856 | 0.02858 | | |
| | -70 | ns | 0.1958 | 0.01248 | 0.008715 | 0.002819 | | | | -70 | ns | 0.2945 | -0.002649 | 0.008539 | 0.01016 | | |
| | -60 | ** | 0.0013 | 0.2311 | 0.1206 | 0.02943 | | | | -60 | ns | 0.6378 | 0.1268 | 0.1131 | 0.02867 | | |
| | -50 | *** | 0.0007 | 0.6187 | 0.2775 | 0.08473 | | | | -50 | ns | 0.4275 | 0.2727 | 0.229 | 0.05384 | | |
| | -40 | ** | 0.003 | 1.147 | 0.5358 | 0.1824 | | | | -40 | ns | 0.3033 | 0.4469 | 0.3571 | 0.08468 | | |
| | -30 | ** | 0.0073 | 1.91 | 0.9128 | 0.3362 | | | | -30 | ns | 0.2427 | 0.6508 | 0.5081 | 0.1177 | | |
| | -20 | * | 0.0157 | 2.7 | 1.387 | 0.5002 | 4; 3 | 14; 10 | | -20 | ns | 0.1948 | 1.049 | 0.8112 | 0.1751 | 2; 2 | 11; 10 |
| | -10 | * | 0.0405 | 3.426 | 1.944 | 0.6806 | | | | -10 | ns | 0.0834 | 1.45 | 1.083 | 0.1978 | | |
| | 0 | ns | 0.0516 | 4.11 | 2.409 | 0.8261 | | | | 0 | ns | 0.1151 | 1.759 | 1.374 | 0.2304 | | |
| | 10 | ns | 0.0661 | 4.826 | 2.919 | 0.9849 | | | | 10 | ns | 0.0907 | 2.051 | 1.59 | 0.2552 | | |
| | 20 | ns | 0.0833 | 5.547 | 3.458 | 1.15 | | | | 20 | ns | 0.083 | 2.334 | 1.814 | 0.2802 | | |
| | 30 | ns | 0.1095 | 6.275 | 4.034 | 1.34 | | | | 30 | ns | 0.0828 | 2.608 | 2.038 | 0.3075 | | |
| | 40 | ns | 0.1237 | 7.01 | 4.598 | 1.502 | | | | 40 | ns | 0.1195 | 2.88 | 2.307 | 0.3495 | | |
| Control vs. Pain/ IgG4 | -80 | ns | 0.8957 | -0.1188 | -0.1146 | 0.03171 | | | Control vs. Pain/ IgG4 | -80 | ns | 0.1605 | -0.1073 | -0.07366 | 0.02281 | | |
| | -70 | ns | 0.2981 | 0.01248 | 0.01973 | 0.006694 | | | | -70 | ns | 0.2159 | -0.002649 | 0.01071 | 0.01017 | | |
| | -60 | ns | 0.1549 | 0.2311 | 0.186 | 0.03053 | | | | -60 | ns | 0.5771 | 0.1268 | 0.1126 | 0.0249 | | |
| | -50 | ** | 0.0054 | 0.6187 | 0.3618 | 0.08131 | | | | -50 | ns | 0.2914 | 0.2727 | 0.2223 | 0.04599 | | |
| | -40 | ** | 0.0043 | 1.147 | 0.6221 | 0.1588 | | | | -40 | ns | 0.1884 | 0.4469 | 0.3433 | 0.07454 | | |
| | -30 | ** | 0.0029 | 1.91 | 0.8865 | 0.2896 | | | | -30 | ns | 0.172 | 0.6508 | 0.4946 | 0.1082 | | |
| | -20 | ** | 0.0036 | 2.7 | 1.232 | 0.4274 | 4; 3 | 14; 12 | | -20 | ns | 0.315 | 1.049 | 0.8525 | 0.1899 | 2; 2 | 11; 11 |
| | -10 | ** | 0.0044 | 3.426 | 1.621 | 0.5438 | | | | -10 | ns | 0.1367 | 1.45 | 1.093 | 0.2298 | | |
| | 0 | ** | 0.0039 | 4.11 | 1.948 | 0.6406 | | | | 0 | ns | 0.1718 | 1.759 | 1.373 | 0.2718 | | |
| | 10 | ** | 0.0035 | 4.826 | 2.278 | 0.7435 | | | | 10 | ns | 0.1885 | 2.051 | 1.624 | 0.314 | | |
| | 20 | ** | 0.0035 | 5.547 | 2.609 | 0.8584 | | | | 20 | ns | 0.2293 | 2.334 | 1.897 | 0.3526 | | |
| | 30 | ** | 0.0034 | 6.275 | 2.946 | 0.97 | | | | 30 | ns | 0.2442 | 2.608 | 2.138 | 0.3914 | | |
| | 40 | ** | 0.0033 | 7.01 | 3.271 | 1.085 | | | | 40 | ns | 0.3332 | 2.88 | 2.444 | 0.439 | | |
| Control vs. Pain/ IgGX | -80 | ns | 0.5019 | -0.1188 | -0.1425 | 0.03484 | | | Control vs. Pain/ IgGX | -80 | ns | 0.6559 | -0.1073 | -0.09375 | 0.03002 | | |
| | -70 | ns | 0.3757 | 0.01248 | 0.0101 | 0.002629 | | | | -70 | ns | 0.1433 | -0.002649 | 0.0133 | 0.01009 | | |
| | -60 | ns | 0.4249 | 0.2311 | 0.1994 | 0.03893 | | | | -60 | ns | 0.6965 | 0.1268 | 0.14 | 0.03328 | | |
| | -50 | ns | 0.1068 | 0.6187 | 0.4452 | 0.1033 | | | | -50 | ns | 0.9553 | 0.2727 | 0.2765 | 0.06621 | | |
| | -40 | ns | 0.0881 | 1.147 | 0.7659 | 0.2135 | | | | -40 | ns | 0.845 | 0.4469 | 0.4258 | 0.1064 | | |
| | -30 | ns | 0.1069 | 1.91 | 1.212 | 0.4153 | | | | -30 | ns | 0.7705 | 0.6508 | 0.6065 | 0.1497 | | |
| | -20 | ns | 0.0971 | 2.7 | 1.696 | 0.5803 | 4; 3 | 14; 11 | | -20 | ns | 0.5231 | 1.049 | 0.9054 | 0.2206 | 2; 2 | 11; 11 |
| | -10 | ns | 0.0963 | 3.426 | 2.186 | 0.7148 | | | | -10 | ns | 0.585 | 1.45 | 1.309 | 0.2534 | | |
| | 0 | ns | 0.0983 | 4.11 | 2.646 | 0.8498 | | | | 0 | ns | 0.6605 | 1.759 | 1.628 | 0.2937 | | |
| | 10 | ns | 0.0943 | 4.826 | 3.088 | 0.9954 | | | | 10 | ns | 0.9562 | 2.051 | 2.032 | 0.3451 | | |
| | 20 | ns | 0.0965 | 5.547 | 3.544 | 1.155 | | | | 20 | ns | 0.9165 | 2.334 | 2.375 | 0.3819 | | |
| | 30 | ns | 0.0913 | 6.275 | 3.982 | 1.3 | | | | 30 | ns | 0.7227 | 2.608 | 2.761 | 0.4246 | | |
| | 40 | ns | 0.0917 | 7.01 | 4.453 | 1.451 | | | | 40 | ns | 0.6225 | 2.88 | 3.113 | 0.4664 | | |
| Control vs. Pain/ No IgG4 | -80 | ns | 0.5248 | -0.1202 | -0.1045 | 0.02372 | | | Control vs. Pain/ No IgG4 | -80 | ns | 0.3863 | -0.1073 | -0.08355 | 0.02681 | | |
| | -70 | ns | 0.1299 | 0.0111 | 0.0147 | 0.00216 | | | | -70 | ns | 0.1801 | -0.002649 | 0.01213 | 0.01038 | | |
| | -60 | ns | 0.427 | 0.1711 | 0.152 | 0.02294 | | | | -60 | ns | 0.8401 | 0.1268 | 0.1206 | 0.03031 | | |
| | -50 | ns | 0.485 | 0.3392 | 0.3047 | 0.04745 | | | | -50 | ns | 0.5046 | 0.2727 | 0.2339 | 0.05708 | | |
| | -40 | ns | 0.41 | 0.5291 | 0.4683 | 0.07036 | | | | -40 | ns | 0.3892 | 0.4469 | 0.3654 | 0.09245 | | |
| | -30 | ns | 0.4563 | 0.7413 | 0.6695 | 0.09215 | | | | -30 | ns | 0.5207 | 0.6508 | 0.5592 | 0.14 | | |
| | -20 | ns | 0.4514 | 1.082 | 0.9609 | 0.154 | 4; 2 | 14; 10 | | -20 | ns | 0.2987 | 1.049 | 0.8366 | 0.1984 | 2; 2 | 11; 10 |
| | -10 | ns | 0.8654 | 1.367 | 1.329 | 0.2201 | | | | -10 | ns | 0.1499 | 1.45 | 1.103 | 0.231 | | |
| | 0 | ns | 0.7912 | 1.674 | 1.605 | 0.2536 | | | | 0 | ns | 0.1612 | 1.759 | 1.349 | 0.2813 | | |
| | 10 | ns | 0.7802 | 1.974 | 1.892 | 0.2848 | | | | 10 | ns | 0.1898 | 2.051 | 1.616 | 0.3201 | | |
| | 20 | ns | 0.8039 | 2.299 | 2.217 | 0.3177 | | | | 20 | ns | 0.2115 | 2.334 | 1.867 | 0.3613 | | |
| | 30 | ns | 0.798 | 2.614 | 2.52 | 0.355 | | | | 30 | ns | 0.2458 | 2.608 | 2.125 | 0.4031 | | |
| | 40 | ns | 0.7058 | 2.961 | 2.813 | 0.3792 | | | | 40 | ns | 0.261 | 2.88 | 2.361 | 0.4477 | | |

aAb=autoantibody presence; Res.=rescue condition; C=control; sub=patient serum subclass; ns=Non-significant; *p<0.05, **p<0.01, ***p<0.0001; N=number of experiments; n=number of cells

**Supplemental Table 5: Potassium channel activity of DRG neurons in the presenvce and absence of toxines**

| | Voltage | Significance | *p*-Value | Mean Ctrl | Mean Toxin | *N* | *n* | | Voltage | Significance | *p*-Value | Mean Ctrl | Mean Toxin | *N* | *n* |
|---|---|---|---|---|---|---|---|---|---|---|---|---|---|---|---|
| Control vs. Conotoxin κM-RIIIJ | -80 | ns | 0.9966 | -0.08765 | -0.14 | 2 | 11 | Control vs.α-Dendrotoxin | -80 | ns | 0.9939 | -0.1249 | -0.1829 | 2 | 10 |
| | -70 | ns | >0.9999 | 0.007773 | 0.005915 | 2 | 11 | | -70 | ns | >0.9999 | 0.00054 | -0.0005738 | 2 | 10 |
| | -60 | ns | >0.9999 | 0.1373 | 0.1709 | 2 | 11 | | -60 | ns | >0.9999 | 0.1587 | 0.1911 | 2 | 10 |
| | -50 | ns | 0.9678 | 0.2816 | 0.3492 | 2 | 11 | | -50 | ns | 0.9993 | 0.3382 | 0.3845 | 2 | 10 |
| | -40 | ns | 0.9375 | 0.4448 | 0.5187 | 2 | 11 | | -40 | ns | 0.9996 | 0.5437 | 0.588 | 2 | 10 |
| | -30 | ns | 0.9997 | 0.6624 | 0.7039 | 2 | 11 | | -30 | ns | >0.9999 | 0.7842 | 0.7807 | 2 | 10 |
| | -20 | * | 0.0172 | 1.145 | 0.9601 | 2 | 11 | | -20 | ns | 0.2614 | 1.16 | 1.025 | 2 | 10 |
| | -10 | **** | <0.0001 | 1.53 | 1.264 | 2 | 11 | | -10 | ** | 0.0074 | 1.538 | 1.33 | 2 | 10 |
| | 0 | **** | <0.0001 | 1.888 | 1.519 | 2 | 11 | | 0 | *** | 0.0003 | 1.912 | 1.651 | 2 | 10 |
| | 10 | **** | <0.0001 | 2.214 | 1.775 | 2 | 11 | | 10 | **** | <0.0001 | 2.34 | 1.937 | 2 | 10 |
| | 20 | **** | <0.0001 | 2.532 | 2.023 | 2 | 11 | | 20 | **** | <0.0001 | 2.748 | 2.225 | 2 | 10 |
| | 30 | **** | <0.0001 | 2.841 | 2.26 | 2 | 11 | | 30 | **** | <0.0001 | 3.121 | 2.509 | 2 | 10 |
| | 40 | **** | <0.0001 | 3.158 | 2.514 | 2 | 11 | | 40 | **** | <0.0001 | 3.485 | 2.787 | 2 | 10 |

*N*=number of experiments; *n*=number of cells recorded; ns=non-significant, *$p$<0.05, **$p$<0.01, ***$p$<0.001.****$p$<0.0001

#### SUPPLEMENTAL FIGURES

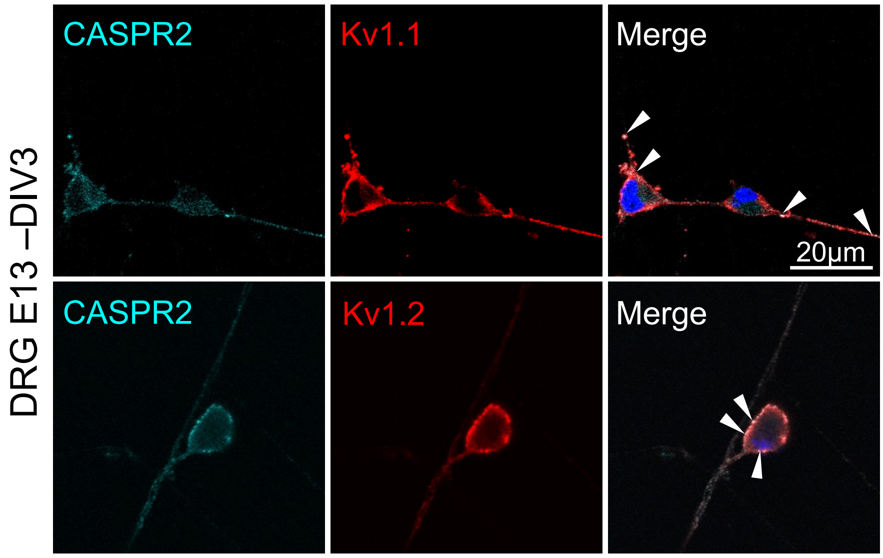


**Supplemental Figure 1**

**Embryonic DRG neurons express CASPR2 and potassium channel subunits.** Embryonic DRG neurons seeded at day 13 (E13) and stained after three days in culture (DIV3) for CASPR2 (cyan) and either the potassium channel subunit Kv1.1 or Kv1.2 (red), DAPI (blue). Colocalized signals (white) of CASPR2 and either Kv1.1 or Kv1.2 are observed at the cellular membrane (white arrow heads).

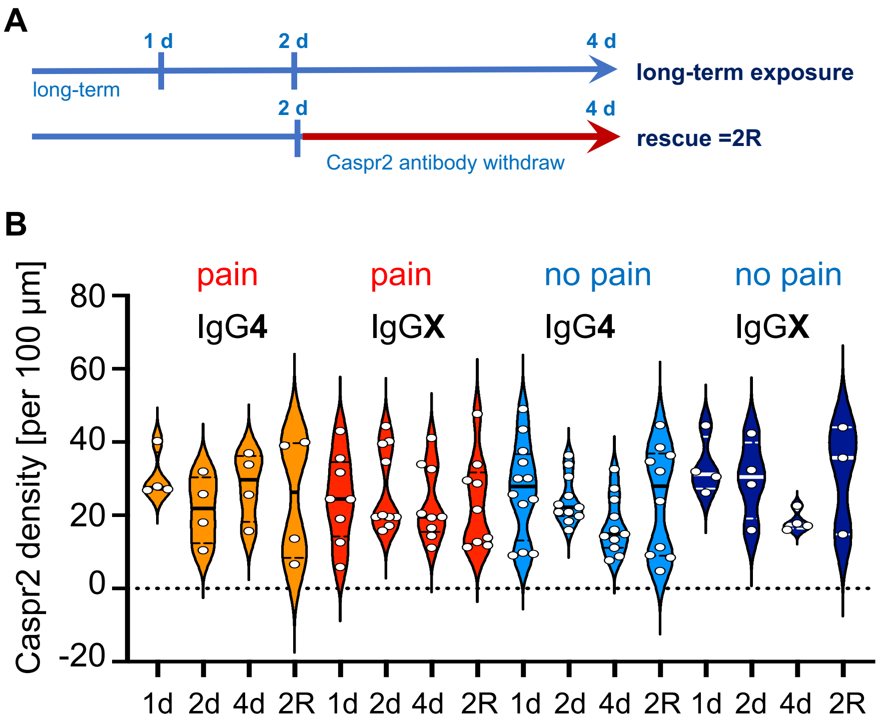


**Supplemental Figure 2**

**CASPR2 expression is unaltered in the presence of different subclasses of CASPR2 autoantibodies.** A) Timeline of the experiments with presence of CASPR2 aAbs for one day (1d), two days (2d), four days (4d), or incubated for 2d followed by a recovery period of additional 2d (=2R). B) CASPR2 aAbs of different subclassifications according to pain/no pain and IgG subclass (IgG**4** only and IgG4 and at least one additional IgG = IgG**X**) were incubated with DRG neurons. Quantification of relative CASPR2 expression density per 100µm axon for four patient serum pools with or without pain and with only IgG4 or additional other IgG subclasses. Data are shown as violin plots with individual values, median=bold line, quartiles=dotted lines. Note, there was always a slight reduction of CASPR2 expression after 4d presence of the aAbs which was rescued upon aAb withdraw.

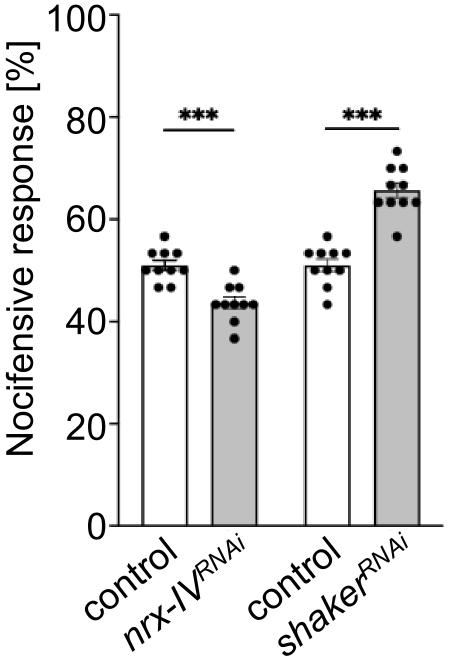


**Supplemental Figure 3: Nocifensive behavior of *Drosophila*.** Knockdown of *neurexin-IV* (*nrx-IV*) specifically in nociceptors (*ppk-GAL4>UAS-nrx-IV*$^{RNAi}$) reduced nocifensive responses of larvae to a mechanical stimulus compared to the genetic control (*ppk-GAL4/+*). In contrast, *shaker* knockdown (*ppk-GAL4>UAS-shaker*$^{RNAi}$) increased nocifensive behavior. Data are presented as mean ± SEM. ****p*≤0.001.

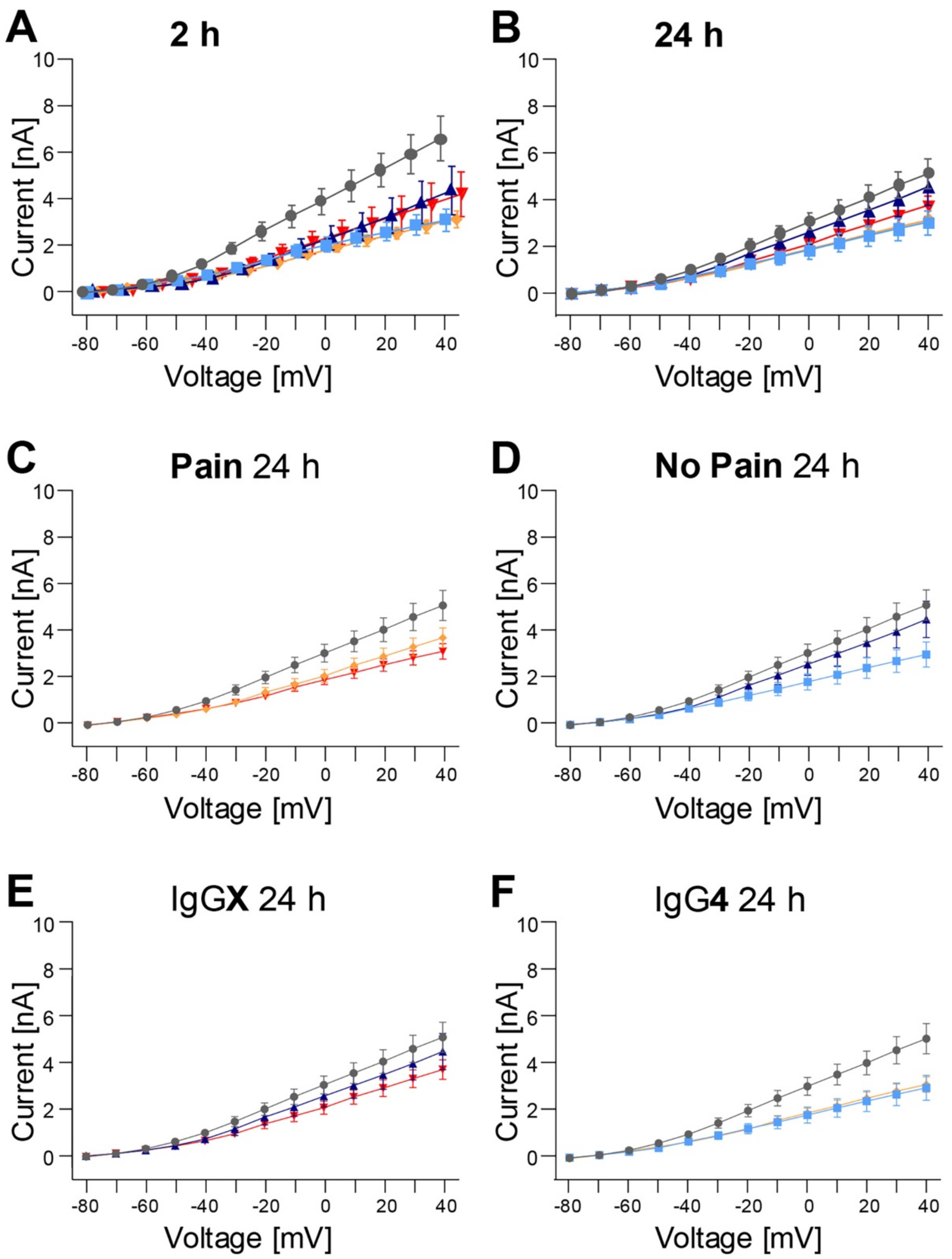


**Supplemental Figure 4**

**Decrease of Kv currents upon 24h presence of CASPR2 aAbs.** A-B) Current-Voltage relation (I-V) plot measured after 2h and 24h aAb application on DRG neurons with healthy control serum (grey), CASPR2 subclassifications: pain IgG4 (yellow), pain IgGX (red), no pain IgG4 (light blue), no pain IgGX (dark blue). C-F) same as in B but subdivided according to pain/no pain phenotype or IgG subclass. Data are presented as mean ± SEM.

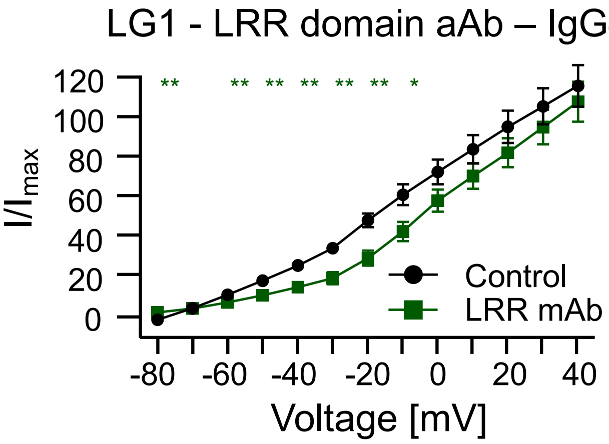


**Supplemental Figure 5**

**Decrease of Kv currents in the presence of a LG1 aAb.** Current-Voltage relation (I-V) plot (voltage step protocol from -80 to +40 mV) measured after 2h aAb application on DRG neurons with healthy control serum (grey), monoclonal LG1 autoantibody targeting the LRR domain of LG1. Data are presented as mean ± SEM. Level of significance: *$p$<0.05, **$p$<0.01.
